# T-cell commitment inheritance – an agent-based multi-scale model

**DOI:** 10.1101/2023.10.18.562905

**Authors:** Emil Andersson, Ellen V. Rothenberg, Carsten Peterson, Victor Olariu

## Abstract

T-cell development provides an excellent model system for studying lineage commitment from a multipotent progenitor. The intrathymic development process has been thoroughly studied. The molecular circuitry controlling it has been dissected and the necessary steps like programmed shut off of progenitor genes and T-cell genes upregulation have been revealed. However, the exact timing between decision-making and commitment stage remains unexplored. To this end, we implemented an agent-based multi-scale model to investigate inheritance in early T-cell development. Treating each cell as an agent provides a powerful tool as it tracks each individual cell of a simulated T-cell colony, enabling the construction of lineage trees. Based on the lineage trees, we introduce the concept of the last common ancestors (LCA) of committed cells and analyse their relations, both at single-cell level and population level. In addition to simulating wild-type development, we also conduct knockdown analysis. Our simulations showed that the commitment is a three-step process over several cell generations where a cell is first prepared by a transcriptional switch. This is followed by the loss of the Bcl11b-opposing function two to three generations later which is when the decision to commit is taken. Finally, after another one to two generations, the cell becomes committed by transitioning to the DN2b state. Our results showed that there is inheritance in the commitment mechanism.

## Introduction

Commitment of T-cell progenitors offers an opportunity to dissect the molecular circuitry establishing cell identity in response to environmental signals. This intrathymic development process encompasses programmed shutoff of progenitor genes^1^, upregulation of T-cell specification genes, proliferation, and ultimately commitment^2^. The core gene regulatory network controlling this process was identified from results from perturbation studies^3,4^. It was also shown that Bcl11b expression correlates with the functional committed state cells and can thus serve as a proxy for commitment^5,6^.

The stages of early T-cell development are well known. The T-cell progenitor cells are CD4 and CD8 double negative (DN) and do not express T-cell receptors. The Kit^high^ early thymic progenitors (ETPs or DN1s) transition to the DN2a state which is marked by CD25 surface expression^7,8^. The DN2a cells upregulate the expression of Bcl11b, which correlates with the commitment to the T-cell lineage fate, as they transition to the DN2b state^5,7–10^. The cells continue through the DN3 and DN4 stages and progress with the late T-cell development, however, these stages of development are not within the scope of this work. The cells proliferate during the DN1 stage with an increasing proliferation rate as the cells progress to DN2. The steps of the early T-cell development are summarised in Fig. 1a and we refer to references^1,2^ for detailed reviews.

**Figure 1.**
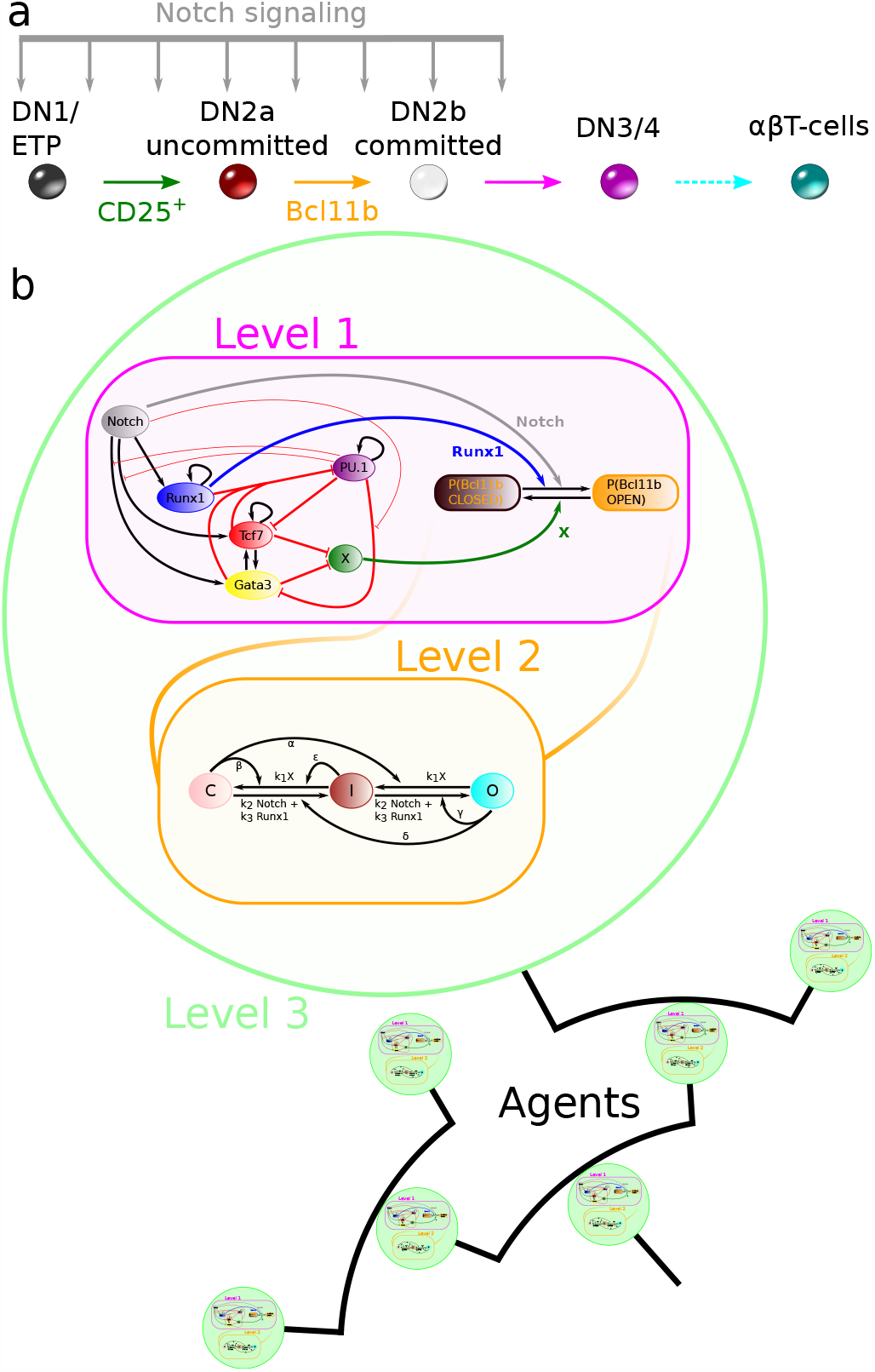
Agent-based multi-scale model for early T-cell development commitment. **a**) Schematics over the early T-cell development. Early thymic progenitors (ETPs or DN1) transition to the DN2a state which is marked by CD25 surface expression. Commitment to the T-cell fate is observed by Bcl11b upregulation as the cells progress to the DN2b state. The T-cell lineage development continues through DN3 and DN4 stages and eventually becomes mature T-cells. The early T-cell development takes place under the influence of Notch signalling. **b**) Depiction of the multi-scale agent-based model. The magenta-coloured box illustrates level 1 and contains the GRN topology. The black arrows and thick red blunted arrows represent positive and negative direct regulation respectively. The thin red blunted arrows represent inhibition of regulation. The blue and grey arrows represent that Runx1 and Notch promote the opening of Bcl11b regulatory sites, while the green arrow shows that X keeps the sites closed. The orange box illustrates the epigenetic mechanism of level 2. The regulatory sites can change between three different states (closed, intermediate and open) and are affected by input signals from level 1. Each cell of level 3 (green circles) contains a copy of levels 1 and 2. The agent-based model implementation tracks the relation between the proliferating cells in lineage trees.

We have earlier developed and independently verified a core gene regulatory network (GRN) governing the early stages of T-cell progenitor commitment^11^. This GRN is built on experimentally shown interactions between T-cell specification genes Tcf7 (TCF1)^5,12^, Gata3^5,13^ and Runx1^4,5^, and the opposing Spi1 (PU.1) gene^3^. The dynamics governed by this network are influenced by an extrinsic T-cell positive input from Notch signalling present due to the thymic microenvironment. It has been shown that Runx1 control the timing of T-cell development and that Tcf7 is needed in order for Bcll1b to turn on^14^. Connecting Bcl11b directly to the core GRN would predict an upregulation of Bcl11b synchronised with the expression of the T-cell-specific factors. It has been experimentally shown that Bcl11b is turned on a few cell cycles after CD25 expression and T-cell-specific factors upregulation^5,15,16^. Therefore, we proposed a model for a transcription level of regulation which propagates into an epigenetic level of Bcl11b regulation^11^. This single-cell model was trained with data from RNA fluorescence in situ hybridization (FISH). Furthermore, the ultimate activation of Bcl11b was found to depend on a slow cis-acting epigenetic mechanism, because direct observation of live cells showed that the two equivalent alleles within a single cell could be activated days apart^17^. Moreover, this two-level regulation model was placed in a proliferation level, resulting in a multi-scale model which allowed us to investigate the T-cell commitment mechanism at bulk level. The predictions from this model were validated with clonal kinetic data. Our multi-scale model predicted that DN1 population developmental heterogeneity can arise solely from GRN noise. It also showed that the observed heterogeneous delay in lineage commitment marked by Bcl11b arises both from GRN and epigenetic stochasticity. Furthermore, we observed that the delay between the loss of the Bcl11b opposing function X and expression of Bcl11b was similar to the delay between the CD25 surface expression and the expression of Bcl11b. However, we could not specify the exact timing of the decision to commit with respect to the observed Bcl11b upregulation or whether there is an inheritance of progression towards the T-cell fate.

To this end, we further develop our model by putting the full multi-scale model in an agent-based setting. By this augmentation, we unlock the ability to study *in silico* single-cell properties of T-cell development. With this framework, we can investigate if inheritance plays a role in the T-cell commitment mechanism. We do so by studying lineage trees of simulated T-cell colonies and introducing the concept of last common ancestors (LCAs) of Bcl11b-positive cells. We find that inheritance does play an important role and that the commitment mechanism takes place over several cell generations. Importantly, we find that a cell’s decision to commit is actually made one to two generations before Bcl11b is upregulated.

## Results

### The model

We investigate inheritance in T-cell commitment by performing *in silico* experiments. We propose an agent-based model with a core consisting of our previously published stochastic multi-scale model^11^. By treating single cells as separate agents, we can construct lineage trees corresponding to an entire cell colony, enabling us to access details like inheritance which was previously not readily available. The model consists of three levels: level 1 is a transcriptional level inside a cell, containing a GRN governing the core differentiation process; level 2 is an epigenetic level, also inside a cell, where the Bcl11b regulatory region is treated by opening and closing of regulatory sites depending on signals from the transcriptional level; and finally, level 3, a proliferation level, where the cells as separate agents divide and pass down properties to their descendants (see Fig. 1b). The mathematical details of all three levels in the agent-based model are described in Methods.

#### Level 1: transcriptional level

The gene regulatory network in the transcriptional level consists of 6 interacting genes: Runx1, Gata3, PU.1 (Spi1), Tcf7 (TCF1), Notch signalling, and a Bcl11b inhibitory function X (see Fig. 1b). X includes a slow initial chromatin opening mechanism and other possible Bcl11b-antagonists, inhibiting the opening of the Bcl11b regulatory sites. Runx1 and Notch act positively for opening the Bcl11b regulatory sites and Notch is promoting Runx1, Tcf7 and Gata3. Both Tcf7 and Gata3 operate towards opening the Bcl11b regulatory sites by inhibiting X and PU.1. In turn, PU.1 inhibits Tcf7 and Gata3 as well as their Notch activation. The stochastic implementation of the GRN model is done with the Gillespie algorithm^18^ (see Methods). The communications between the transcriptional level and epigenetic level are conducted through the signals from X, Runx1 and Notch.

#### Level 2: epigenetic level

Bcl11b becomes expressed in the transition from the T-cell development stage DN2a to DN2b. In our model, Bcl11b is regulated by an epigenetic mechanism consisting of the opening and closing of regulatory sites. The 500 Bcl11b regulatory sites can either be closed (C), intermediate (I) or open (O). Cells at generation 0 have the Bcl11b regulating region completely closed. Cells with more than 75 % of the regulatory sites open are considered to have an open Bcl11b regulatory region. The epigenetic level is regulated by the transcriptional level. Runx1 and Notch increase the probability of opening regulatory sites while X increases the probability of closing them. More details about the stochastic implementation can be found in Methods.

#### Level 3: proliferation level

The proliferation level simulates an entire cell population. The simulations always start with one individual cell at generation 0. The dynamics of the multi-scale model containing the transcriptional and epigenetic levels inside the cell are simulated over time. Cell division is also implemented with the daughter cells containing the same multi-scale model as the mother, continuing the transcriptional and epigenetic simulations. The colony evolves through multiple divisions and its simulation corresponds to 120 h in experimental time. The division times are sampled from distributions that vary with cell generations. This creates heterogeneity of the age of cells from different generations, i.e. one simulated colony undergo 6 cell cycles while another undergoes 11.

During cell division, all the gene expression levels are inherited keeping them at the same levels as inside the mother cell. However, the divisions follow a set of conditional rules imposed on the regulatory sites^19^. If X is present, the regulatory sites are copied from the mother cell, while the loss of X promotes the opening of the regulatory sites. Thus, a strong driving force to opening up the Bcl11b regulatory region is the repression of X.

### Agent-based model

Our agent-based implementation enables us to record the relations between the cells within a lineage tree. This way, we can investigate the existence of inheritance in T-cell commitment. This type of modelling makes it possible to know the transcription levels, epigenetic status and age of each cell at any simulation time point. Having access to this type of cell information is very informative for future experimental efforts.

### Investigating inheritance

As illustrated in Fig. 1a, surface expression of CD25 marks the transition from ETP to DN2a and Bcl11b expression marks the T-cell commitment. Bcl11b heterogeneity was shown in colonies of differentiating T-cell progenitors both *in vitro* and *in silico* in Olariu et al. (2021)^11^. Furthermore, for *in vitro* experiments, a large heterogeneity was observed for the number of cell divisions and the time when the colonies reached 50 % CD25 positive cells and 50 % Bcl11b positive cells.

These findings raise a few questions which we attempt to answer using our new agent-based implementation of the multi-scale model:

- What are the mechanisms leading to the observed heterogeneity within a single colony?
- When is the decision taken to open or keep closed a cell’s Bcl11b regulatory region?
- How are system perturbations affecting the decision to commit?

### Lineage trees

To answer the questions above, we use a tool that represents the simulated colonies as phylogenetic trees^20^ or lineage tree diagrams. Fig. 2a illustrates a simple version of a lineage tree, with key features highlighted. The initial cell at generation 0 is the root of the tree and every cell division leads to two new branches. Each cell is a node in the tree and the radial length of a connection in the tree is proportional to the cell’s lifetime, i.e. the time-axis points radially outwards. The colour of the node represents the status of the cell where black depicts a cell with the Bcl11b regulatory region closed and X expression greater than 0, a red node represents a cell where the Bcl11b is closed but X is not expressed, and white represents a cell where the Bcl11b region is open. With this colouring scheme, one can observe heterogeneity of open Bcll1b within a colony along with the delay between the loss of X and the opening of Bcl11b.

**Figure 2.**
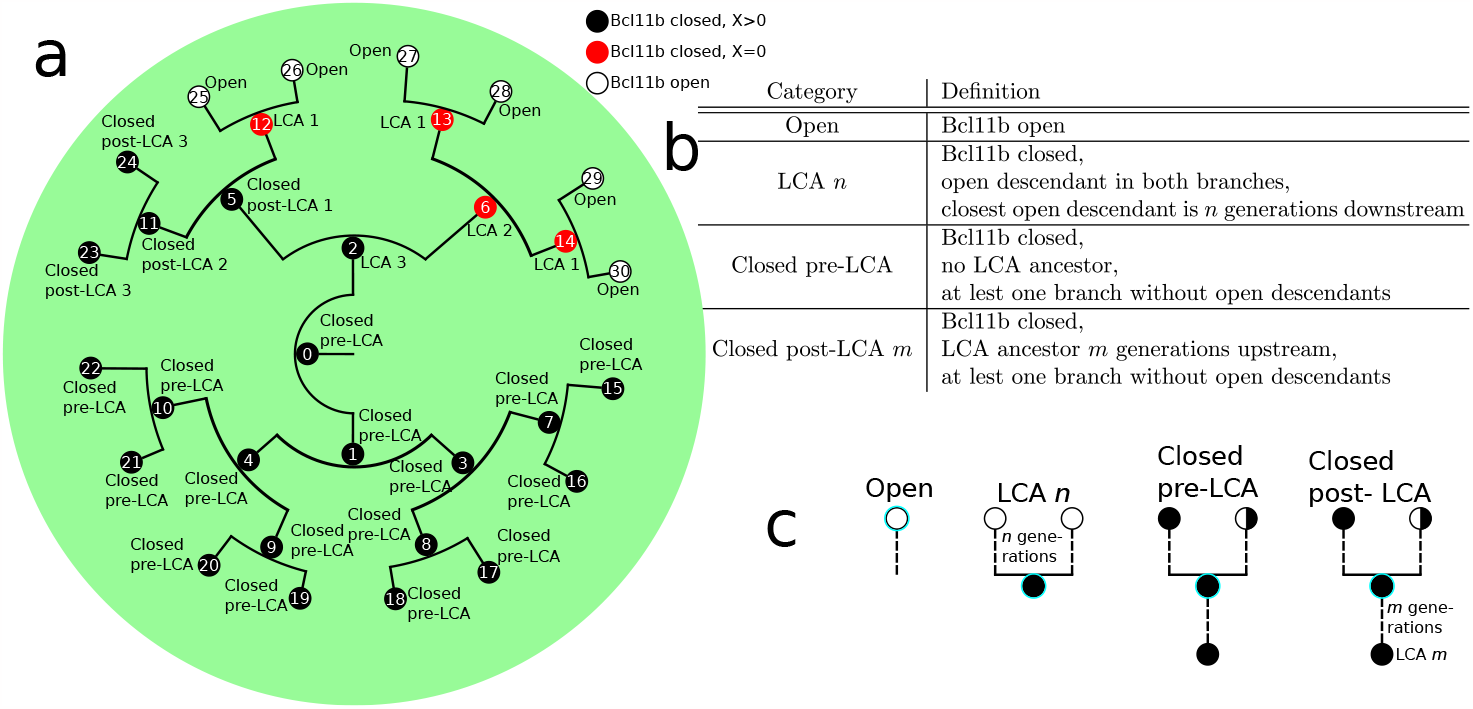
Small lineage tree and last common ancestor (LCA) definitions. **a**) A lineage tree example. Every node is a cell which is uniquely identifiable by an index. Each cell branches into two daughter cells. The radial connecting lines are proportional to a cell’s lifetime. The colour of the node indicates the status of the cell’s Bcl11b regulatory region and X expression, as described by the legend. Each cell is labelled with its LCA label. **b**) Definitions of the LCA categories. **c**) The graphical definition of each LCA category for the cells encircled with cyan. Note the subtle difference between ‘closed post-LCA *m*’ and ‘closed pre-LCA’, i.e. the only difference is whether they have an LCA-ancestor or not.

### Last Common Ancestors

Since the delay between loss of X function along with CD25 activation and Bcl11b opening varies between cells, both in time and number of divisions, we have to track time relative to interesting events rather than experimental time or generation counting^11^. The cells’ main characteristic considered here is whether the Bcl11b regulatory region is open or closed. This is due to the fact that a cell with Bcl11b open has surely committed to the T-cell fate and has become DN2b. Therefore, we introduce a system of categorising the cells uniquely according to their relations with cells with open Bcl11b within a simulated lineage tree (see Figs. 2b and 2c).

We define the first category ‘open’ which includes all cells with open Bcl11b region. Two arbitrarily chosen cells from the ‘open’-category which have mother cells with Bcl11b closed have a last common ancestor (LCA) which is the last Bcl11b-closed cell they both originate from. Therefore, we define any Bcl11b-closed cell that has at least one ‘open’ descendant in each of its two branches as an ‘LCA’-cell. The ‘LCA’-cells are further classified depending on how many generations down the lineage the closest ‘open’ cell is located. For instance, if a closed cell’s both daughter cells are ‘open’, it is of order 1, i.e. categorised as ‘LCA 1’. If a closed cell instead has an ‘open’ cell five generations downstream in one branch, but six generations downstream in the other branch, it is of order 5, i.e. ‘LCA 5’.

Bcl11b-closed cells upstream from the earliest LCA in a lineage are defined as ‘closed pre-LCA’. The so-far unclassified cells are Bcl11b-closed but located downstream from LCAs. These are defined as ‘closed post-LCA’. These cells are also given an order depending on how many generations downstream from the latest LCA the cell is. The definitions of the four categories are summarised in Figs. 2b and 2c.

All the cells in the example tree in Fig. 2a have been categorised according to this system. From this tree, it is clear that a colony can be heterogeneous in Bcl11b. The root cell, cell 0, is ‘Closed pre-LCA’ since every cell in its lower branch, rooted in cell 1, is closed. The top half of the tree, rooted in cell 2, has both open and closed sub-branches. Cell number 2 is an LCA 3’ cell since both its daughters have descendants which are ‘open’, where the closest is three generations away. Cell 2 together with cell 5 (which is a ‘closed post-LCA 1’-cell) are interesting cells since their branches have different fates; some sub-branches open up while others stay closed. Branching points like these are key points to investigate in order to elucidate the inheritance mechanism in T-cell commitment.

By defining these categories, we can examine the internal relations between the cells in a tree in a generation-number-independent way. This is important since some lineage trees may only have 6 generations while others could reach 11. The LCA-category system is a tool to dissect when the commitment decisions happen. Presumably, no decision should have been made before an LCA-cell, thus, all ‘closed pre-LCA’-cells should have similar gene expressions. By indexing the LCA-cells, it is possible to track how many generations before the observed cell state transition the actual decision was taken. One example when the decision to commit is taken very close to the actual cell state transition is if all LCAs of order 2 or higher are similar to ‘closed pre-LCA’. Another very different example is if LCA-cells of order 6 express similar traits as open cells, then the decision to commit is taken long before the observed transition. The ‘closed post-LCA’-cells serve as an indicator for how long it takes for cells that almost made the decision, but did not, to return to the fully closed state.

### *In silico* simulations of lineage trees

We simulated 300 wild-type (WT) T-cell committing colonies, all identically initialised in the ETP cell state, and produced corresponding lineage trees for each colony. Fig. 3a shows an example lineage tree and in Supplementary Figs. 4 to 9 a larger subset is presented. The lineage tree in (Fig. 3a) branched seven times, corresponding to seven divisions, yielding 128 final cells. Out of these, 22 are open, all located at the lower half of the tree. Every cell in the branch rooted at cell 10 is open, and the remaining open cells are all located in the branch rooted at cell 8 (from here, a branch rooted at cell *n* is shorted to branch *n*). The cells in branch 6 show no sign of being about to open up, while a few cells in branch 5 have lost X expression but are still closed.

**Figure 3.**
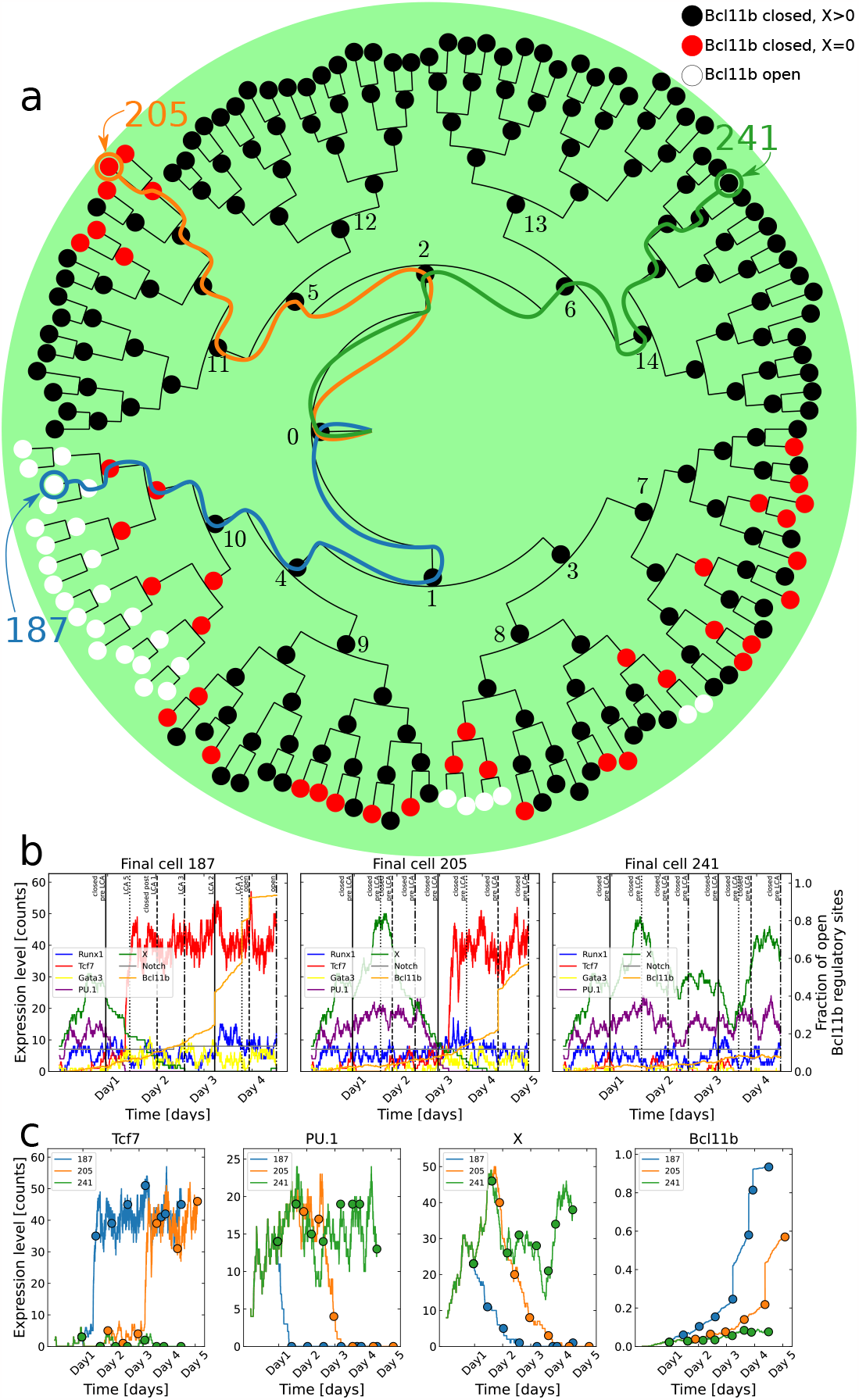
Stochastic simulations of cell lineages. **a**) Lineage tree for a simulated colony. Three different lineages with different fates are marked with blue, orange, and green respectively. Black nodes depict cells with the Bcl11b regulatory region closed and X expression greater than 0, red nodes represent cells where the Bcl11b is closed and X is depleted, and white nodes represent cells where the Bcl11b region is open. **b**) Each panel shows the gene expression dynamics for each of the three marked lineages respectively. The vertical lines represent cell divisions and are labelled with the cells’ corresponding last common ancestor (LCA) categories. **c**) Each panel show the expression for Tcf7, PU.1, X and the fraction of open Bcl11b regulatory sites respectively for the three marked cell lineages. The coloured dots indicate cell divisions.

This tree shows clear heterogeneity in the different branches which makes it a prime candidate to investigate what happens during development at different important branching points. The simulated gene expression levels are shown for three chosen cell lineages in Fig. 3b, where each panel shows the evolution of one lineage. The three lineages are 187 (blue path in Fig. 3a which is an open cell in branch 10; 205 (orange path) which is a closed cell in branch 5, but with X depleted; and 241 (green path) which is a closed cell in branch 6. The vertical lines in Fig. 3b indicate cell divisions and are labelled with the cells’ LCA labels. Fig. 3c shows the expression of Tcf7, PU.1, X and Bcl11b in one panel each to more conveniently compare how these genes differ in the three lineages. Here, the coloured dots indicate cell divisions.

The three lineages only have the first cell in common. It is clear that already after the first division, the blue path starts to become different from the green and orange. PU.1 decreases at the same time as Tcf7 starts to increase. These two components are tightly connected in the GRN (Fig. 1b). PU.1 represses Tc7 both directly and through repression of the Tcf7-activating Notch signal, while Tcf7 jointly represses PU.1 together with Runx1 and Gata3. Runx1 is stable over time and Gata3 follows Tcf7 but is expressed at a much lower level. Therefore, it is probably a small fluctuation of either decreasing PU.1 or increasing Tcf7 which throws the preparatory switch towards the T-cell fate to start producing an excess of Tcf7. Once there is an abundant amount of Tcf7, PU.1 is firmly repressed while Tcf7 is kept at a high level by both its direct self-activation and its indirect activation through Gata3. The orange lineage also goes through this switch, but at a later time point on day 3, while the green lineage does not go through this switch at all. Once Tcf7 (and Gata3) is highly expressed, X is repressed. For the blue lineage, X is depleted before day 3 while, for the orange lineage, this happens at day 4. On the other hand, X stays highly expressed in the green lineage where PU.1 is also high and Tcf7 is low. For the blue and orange lineages, when Tcf7 has gone through the switch and X starts to decrease, the fraction of open Bcl11b regulatory sites slowly starts to increase. However, when X is completely depleted, the division rules promoting the opening of the Bcl11b regulatory sites are kicking in, resulting in Bcl11b making great increasing leaps at every cell division. The blue lineage opens up Bcl11b two generations after X is depleted, exhibiting a delay between the loss of X and the opening of Bcl11b. Presumably, in the orange lineage, Bcl11b would also open up after the next division, if the simulation would have been longer; the orange lineage exhibits similar traits as the blue one but roughly two days delayed. The green lineage is similar to the early behaviour of the blue and orange lineages and would probably also be able to throw the preparatory T-cell fate switch if the simulation was run much longer, this way giving the system a chance to allow high expression of Tcf7 and low PU.1.

### Statistics of multiple simulated cell colonies

In the previous section, we dissected the gene expression dynamics for three specific cell lineages in the lineage tree shown in Fig. 3. In this section, we use the LCA labels (Figs. 2b and 2c) to gauge the commitment dynamics at the population level over all 300 colonies.

The number of divisions in the 300 simulated colonies spanned between five and twelve, with eight or nine divisions being the most common. The colonies were classified according to their LCA properties as described in Sections Last Common Ancestors and Methods and Fig. 6. The mean expression for each gene was calculated for each LCA category, using the cells’ final gene expression values before division. The mean expression levels for Tcf7, PU.1 and X and the fraction of open Bcl11b regulatory sites for the ‘closed pre-LCA’, ‘open’ and ‘LCA *n*’ categories are shown in Fig. 4. The categories are sorted in an approximate developmental order. The grey numbers in each bar indicate the number of cells belonging to each LCA category. The mean expressions for all the genes and for the ‘closed post-LCA *m*’ categories are shown in Supplementary Figs. 1 and 2.

**Figure 4.**
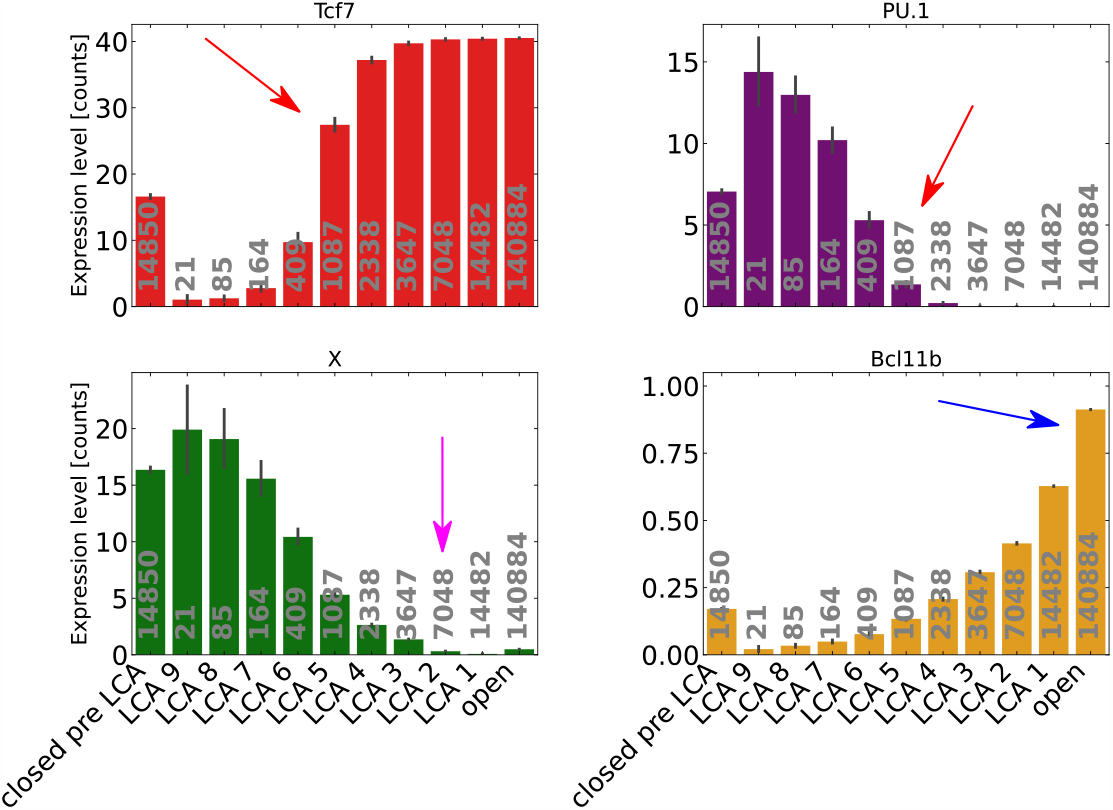
Last common ancestor (LCA) statistics. Mean expression level for Tcf7, PU.1, X, and the fraction of open Bcl11b regulatory sites for cells belonging to the different LCA categories. The categories are ordered in an approximate developmental order. The grey numbers indicate the number of cells belonging to each category. The red arrows point out the preparatory switch towards the T-cell fate. The pink arrow indicates the most common LCA stage where X function is lost. The blue arrow shows that the opening of the Bcl11b regulatory region is delayed with two generations. The error bars represent standard deviations.

Fig. 4 shows development steps from early closed cells to open cells. A first notable feature is that the early LCAs (LCA 9-6) show very little in common with the later LCAs and open cells. It is first at LCA 5 and later where LCA-cells start to have more similar gene expression to the open cells. It is only the Runx1 level that stays fairly stationary through all stages (Supplementary Fig. 1). At LCA 5, the Tcf7 level becomes substantially higher than it is for ‘closed pre-LCA’, while PU.1 becomes substantially lower, as indicated by the red arrows. The high value of Tcf7 and low value of PU.1 ensures that X, after a few generations of GRN simulated dynamics, will become depleted, see the pink arrow at LCA 2. The division rules governed by the absence of X lead to the descendant cells opening up the Bcl11b regions one or two generations after depletion of X at ‘LCA 2 or 1’, see the blue arrow. This chain of events suggests that the ETP cells commit to the T-cell fate several generations before the observed transition to the DN2b state. This result argues for inheritance presence in T-cell commitment. The events highlighted in the example lineages in Section *In silico* simulations of lineage trees and Fig. 3 agree well with the statistics gathered from the 300 simulated lineage trees.

The mean gene expressions for the cells that are closed post-LCA are shown in Supplementary Fig. 2. Note that the ‘closed post-LCA’-cells have two possible future states: they can either continue being closed or become LCA-cells again. Returning to LCA-state is possible if downstream cells in an only-closed-cells branch become open. Supplementary Fig. 2 shows that ‘closed post-LCA 1-3’ cells are all similar to ‘LCA 5-4’ regarding Tcf7 and PU.1 expressions, see the red arrows. These cells can in the following generations either go back to be LCA-cells and eventually open up, or they can continue to stay closed and move to higher ‘closed post-LCA’-orders. Depletion of X by the stochastic process probably governs whether the cells return to the LCA-path, or if they stay closed. The cells that stay closed, i.e. high order of ‘closed post-LCA’, return to a ‘closed pre-LCA’-like state.

### In silico knockdown simulations

We performed *in silicio* knockdown (KD) simulations of Runx1, Tcf7, Gata3 and PU.1 by reducing the production rate and initial expression level of the respective knocked-down gene. Deterministic simulations showed that knocking down the production rates of each gene to 20 % (i.e. 5*×* reduction) perturbs the system sufficiently to result in a dramatic change of the dynamics (Supplementary Fig. 3). For each KD experiment, we simulated 60 colonies each for every KD gene. Figs. 5a and 5b show the average fraction of open cells and the average expression level of X per colony at day 5, respectively. The results in Fig. 5a show that knockdown of PU.1 leads to more Bcl11b-open cells than wild-type (WT) while knockdown of Runx1, Tcf7 and Gata3 results in very few Bcl11-open cells. More specifically, the KD simulation of Tcf7 yielded no open cells while the knockdown of Gata3 led to only one colony with open cells. The average X expression is increased by KD of Runx1, Tcf7 and Gata3, with the greatest effect from Tcf7. KD of PU.1 instead depletes X. Notably, the distributions of the X expression under KD conditions become more narrow compared to the WT, suggesting that the colonies become less heterogeneous during KD.

**Figure 5.**
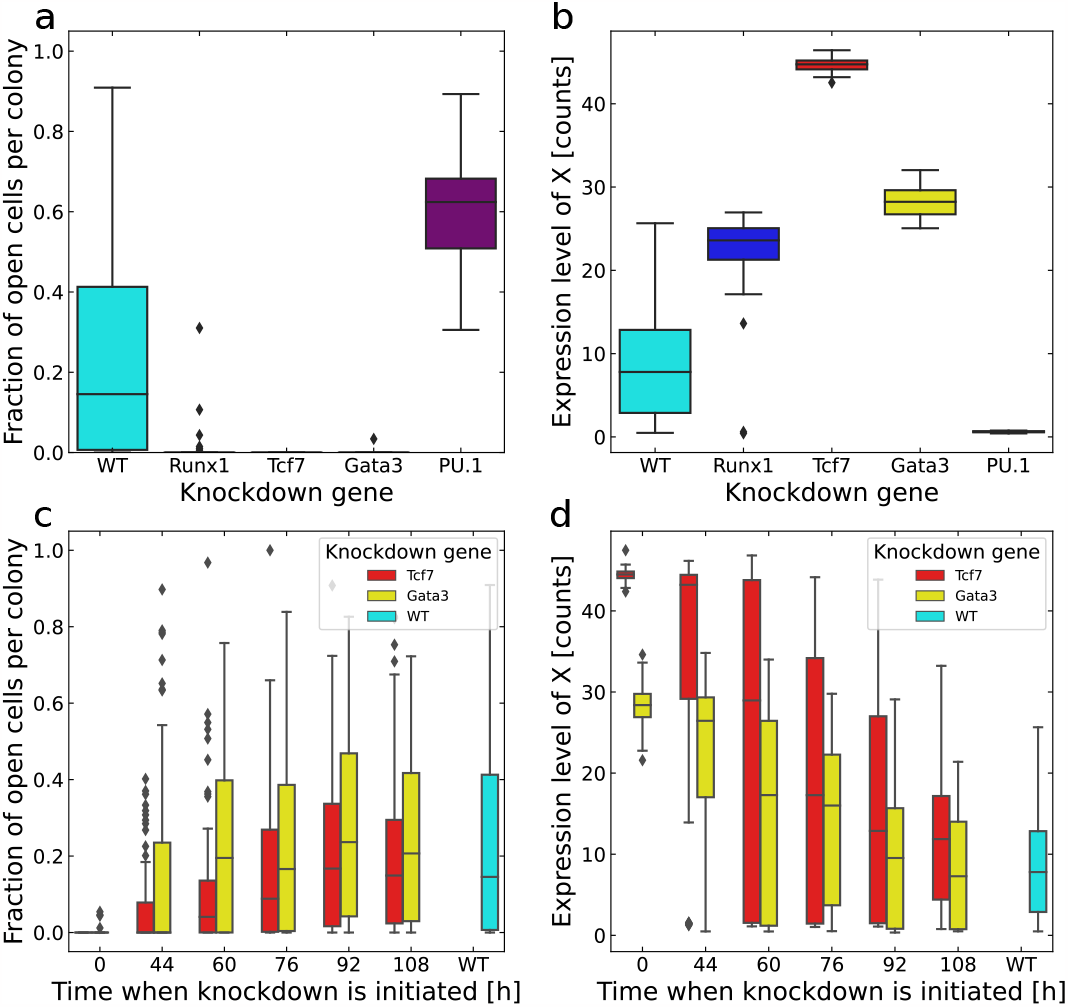
Knockdown simulations. **a**) Statistics on cells with open Bcl11b regulatory regions from KD simulations of Runx1, Tcf7, Gata3 and PU.1 compared to WT simulations where 60 colonies per simulation were considered. Both the genes’ initial transcription counts and production rates are reduced to 20 % of the original values. **b**) Statistics on the expression level of X from the same simulations as in **a**). **c**,**d**) Knockdown simulations of Tcf7 and Gata3 at different time points compared to WT simulations with 180 colonies per simulation type. **c**) shows the distribution of the fraction of cells with open Bcl11b regulatory regions per colony and **d**) shows the expression level of X.

To further dissect the commitment mechanism, we performed simulations where Tcf7 and Gata3 were knocked down at different simulation time points. We simulated KD of the two genes after 0, 44, 60, 76, 92, and 108 hours for 180 colonies for each KD time point and gene. Fig. 5c shows the distribution of the fraction of open cells after 120 hours, across 180 colonies for each KD time considered. The KD of Tcf7 is shown in red and Gata3 in yellow. When KD was initialised from the start, almost none of the colonies had any open cells at the end of the simulation, in accordance with the results shown in Fig. 5a. The fraction of open cells increases with later initialisation of KD where later KD leads to behaviour more similar to WT. The same effect is seen in Fig. 5d for the average expression of X. X is high when Tcf7 or Gata3 KD is conducted early and it decreases with later KD. These results are in accordance with the observed irreversibility of the T-cell commitment process. Previous experimental data showed that Tcf7 and Gata3 are stage-dependent and are more important during the DN1 than the DN2 stage as well as that T-cell commitment is severely hindered by Tcf7 knockdown^5,21^.

### Inheritance investigation results

The observed intra-colony heterogeneity for the Bcl11b regulation site states along with the timing for its opening and closing can be explained by multiple stochastic elements. One important source of stochasticity is the intrinsic noise in the governing GRN. Our simulations showed that the fluctuating expression levels of the transcription factors are responsible for activating the switch towards the T-cell fate. Furthermore, the varying cell cycle length of the cells introduces stochasticity through its effects on the epigenetic level since cell division accelerates the opening of the Bcl11b regulatory sites once X has become depleted. Thus the combined effect of the transcriptional and epigenetic noise and varying cell cycle lengths contributes to the unsynchronised opening of the cells and creates heterogenous cell colonies.

A simulated cell cannot commit to the T-cell lineage without first being prepared by activating the transcriptional switch towards the T-cell fate. However, the decision to open Bcl11b and actually commit to the T-cell fate is taken when the function X is depleted. Until then, the cells can still revert back to their original state as shown by the analysis of the ‘Closed post-LCA’ cells (Supplementary Fig. 2). This is also supported by the knockdown analysis at different time points (Section *In silico* knockdown simulations and Figs. 5c and 5d). The fact that early KD of Tcf7 and Gata3 lead to no or just a few open cells along with high X expression shows that the cells indeed need to be prepared by the thrown transcriptional switch in order to open. Additionally, the late knockdown had no effect compared to WT which shows that Tcf7 and Gata3 become redundant after the cells have committed. Since X expression is kept high after early KD and is lost one or two generations before cells open, we can conclude that the loss of X activity is instrumental for commitment. The KD simulations also show that both Tcf7 and Gata3 are required for the ETP cells to be able to commit to the T-cell fate, even though Tcf7 is more strongly expressed and shows a clearer switch-behaviour (Fig. 3b). Therefore, our results show that there is a one or two-generation delay between the decision to commit and the acquisition of the T-cell committed state, i.e. the Bcl11b-open state.

The LCA analysis showed that not all Bcl11b-closed cells have the same properties. Some closed cells are on the path of becoming Bcl11b-open and are very similar to the open cells in terms of gene expressions. These cells may either be between the switch towards the T-cell fate and X depletion (i.e. between preparation and deciding to commit) or after X depletion but before opening (i.e. between deciding to commit and actually committing). Other closed cells are more similar to the starting ETP cell state. Then the cells are either ‘Closed pre-LCA’ or they can also be ‘Closed post-LCA’. For the latter case, the cells have gone through the preparatory switch but X never became depleted, thus not allowing Bcl11b to become open, resulting in the cells eventually reverting back to a state similar to its initial state. However, gaining back X function after complete depletion is a rare event and it is due to the stochastic implementation of the function X as a transcription production function. As shown in Fig. 4, the statistics over the 300 simulated colonies clearly reveal that X function is most likely lost at the ‘LCA 2’ stage which is roughly two generations before the Bcl11b regulatory region is opened.

## Discussion

In this study, we have used a new agent-based version of our previously published multi-scale model for early T-cell development to explore the role of inheritance in the commitment to the T-cell fate. We developed an analysis tool based on the concept of last common ancestors in phylogenetic analysis^20^ and applied it to T-cell lineage trees. Here, we defined different LCA categories based on the uncommitted cells’ relations to committed DN2b cells. Then we uniquely classified each cell in a simulated lineage tree into the defined categories. We performed single-cell simulations for proliferating wild-type T-cell colonies along with simulations of key gene knockdown. Analysis of both individual colonies as well as statistics for all colonies resulted in an agreeing picture of the T-cell commitment process. The commitment process is a chain of events taking place during multiple cell generations, initiated by the stochastic nature of transcription. The cells first need to be prepared for commitment, which happens if the expression level of PU.1 becomes low, in combination with Tcf7 expression level increase. The governing GRN drives the genes’ expression levels towards this state; however, the analysis of individual cell lineages showed that there is a large heterogeneity in switching to this state. When a cell has been prepared by this switch towards the T-cell fate, the Bcl11b-opposing X function can become depleted. It has been shown both *in vitro* and *in silico*^5,11^ that there is a delay between depleting X (speculated to have the same timing as CD25 upregulation) and Bcl11b-opening. However, the role of X depletion was not completely revealed. In this study, we show *in silico* that depleting X is the decision for a cell to commit to the T-cell fate. Since the observed commitment happens one or two generations after X depletion when a cell opens its Bcl11b-regulatory region and transitions into the DN2b state, we conclude that there is inheritance and it plays an important role in T-cell commitment. Moreover, before X depletion, a cell can still revert to its previous state.

T-cell development has been extensively studied both through experimental and computational efforts. This makes it an excellent model system for studying lineage commitment in other biological systems. This study showing that inheritance plays an important role in T-cell commitment paves the way for revealing whether inheritance is present and is important in the commitment of other cell types. Our modelling framework shows that the epigenetic regulatory level is an important source of delay between commitment decision-making and experimentally observing commitment. Moreover, it shows that the initiation of decision-making is linked to the stochasticity of the transcriptional program. Therefore, transcriptional and epigenetic mechanisms are central to inheritance in cell commitment.

By introducing the LCA category system, we obtained a common framework which allowed us to compare cells in relation to the commitment event instead of absolute time or generation numbers. This enabled us to study and find the steps of the commitment mechanisms at population level. Although every cell can be uniquely classified into the introduced LCA categories, it would be possible to define other sets of categories as well. This should not be considered as a weakness of the method, but rather as a strength since another set of categories could be used to study the mechanism from another angle or other properties. The concept could also be applied to completely different systems with inheritance.

Stochastic elements are integral parts in all the levels of the multi-scale model and can explain the heterogeneity in the timing of the commitment as well as the fraction of committed cells of the simulated *in silico* T-cell colonies. Interestingly, the distribution of the X level becomes more narrow when any of the four genes Runx1, Tcf7, Gata3 and PU.1 is knocked down. From a statistical physics point of view, this corresponds to that the entropy of the cell is decreased when knocking down a gene. The entropy corresponds to a cell’s developmental potential, where high entropy means that the cell is at a branching point in the developmental path with many options^22^. Thus, knocking down a gene makes the population more homogeneous which decreases the entropy and closes off potential developmental branches. This forces cells to either commit to the T-cell fate as for PU.1 KD or leave the T-cell path as for the other KD genes.

In summary, our agent-based multi-scale model predicts that inheritance of progression towards T-cell fate plays a significant role in the commitment mechanism. Moreover, the model shows that the commitment takes place over several cell generations. The most striking model prediction is that a cell’s decision to commit is actually made one to two generations before observed commitment, i.e. Bcl11b upregulation. Since the T-cell development represents an important model system, these results carry relevance for development in other biological systems.

## Methods

The agent-based multi-scale model was implemented from scratch using Python 3.7^23^. The code is available at https://github.com/Emil-cbbp/agent-based_multi-scale_model.git. All the tuning of the model was done in our previous study^11^ and we left the parameter values unchanged. For the independence of this article, we summarise the multi-scale model in the following section.

### Model implementation

#### Level 1: transcriptional level

The GRN in the transcriptional level was modelled using a set of rate equations following the Shea-Ackers formalism^24^. The GRN includes Runx1, Tcf7, Gata3, PU.1, X and Notch signalling where the concentration levels of these are denoted [*R*], [*T*], [*G*], [*P*] and [*X*] respectively. Notch signalling is denoted with *N*. The rate equations used are

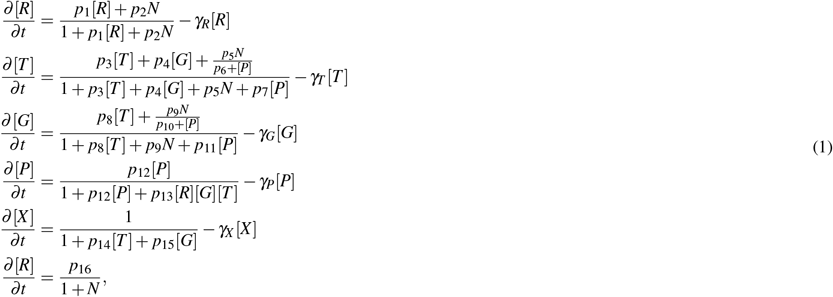

with parameter values listed in Table 1. We refer to Olariu *et al*. (2021)^11^ for a detailed motivation of the rate equation construction and parameter optimisation. The simulations were conducted using the stochastic Gillespie algorithm^18^ implemented from scratch.

**Table 1.** Parameter values for GRN in the transcriptional level.

| Parameter | Value |
| --- | --- |
| $p_1$ | 0.10 |
| $p_2$ | 1.00 |
| $p_3$ | 5.00 |
| $p_4$ | 1.00 |
| $p_5$ | 1.50 |
| $p_6$ | 0.01 |
| $p_7$ | 0.50 |
| $p_8$ | 0.70 |
| $p_9$ | 0.50 |
| $p_{10}$ | 1.00 |
| $p_{11}$ | 0.20 |
| $p_{12}$ | 2.50 |
| $p_{13}$ | 2.60 |
| $p_{14}$ | 2.00 |
| $p_{15}$ | 1.00 |
| $p_{16}$ | 0.01 |
| $\gamma_R$ | $0.15 \text{ h}^{-1}$ |
| $\gamma_T$ | $0.15 \text{ h}^{-1}$ |
| $\gamma_G$ | $0.23 \text{ h}^{-1}$ |
| $\gamma_P$ | $0.06 \text{ h}^{-1}$ |
| $\gamma_X$ | $0.02 \text{ h}^{-1}$ |

#### Level 2: epigenetic level

We used an epigenetic model controlling the Bcl11b regulation adopted from Haerter *et al*. (2014)^19^ and Olariu *et al*. (2016)^25^. The model consists of 500 regulatory sites where each can be in one of three states: closed (*C*), intermediate (*I*) or open (*O*). The Bcl11b region is considered to be open if 75 % of the sites or more are open. The epigenetic level receives input signals from the transcriptional level where Runx1 and Notch signalling work toward opening the regulatory sites and X closes them. The states of the regulatory sites are controlled by a set of rate equations,

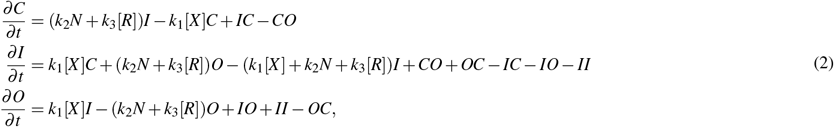

where *C, I* and *O* denotes the number of sites in the respective state (see left half of Table 2 for the parameter values^11^). Not all transitions between states are possible, a site cannot transition directly between *O* and *C* without going through *I*. Some transitions can only take place by the help of a mediator site with a certain state. The right part of Table 2 lists all the possible transitions. The simulations were performed stochastically with an extended version of the Gillespie algorithm^18^. The first extension is that the number of regulatory sites is constant, so when a site transitions from e.g. *O→ I* 1 is added to *I* and subtracted from *O*. Next, when a transition has been chosen, a specific regulatory site is chosen at random. If the site’s state does not match the chosen transition, nothing happens. If one of the five transitions that require mediation is chosen, an additional mediator site is chosen by random. That site also needs to match in order for the transition to take place.

**Table 2.** Parameter values for epigenetic model.

| Parameter | Value | Probability | Transition |
| --- | --- | --- | --- |
| $k_1$ | 0.28 | $k_1X$ | $O \rightarrow I$ |
| | | $k_1X$ | $I \rightarrow C$ |
| $k_2$ | 0.20 | $k_2N + k_3R$ | $C \rightarrow I$ |
| $k_3$ | 0.20 | $k_2N + k_3R$ | $I \rightarrow O$ |
| $\alpha$ | 0.002 | $\alpha$ | $O \rightarrow I$ if mediated by $C$ |
| $\beta$ | 0.002 | $\beta$ | $I \rightarrow C$ if mediated by $C$ |
| $\gamma$ | 0.0005 | $\gamma$ | $I \rightarrow O$ if mediated by $O$ |
| $\delta$ | 0.0005 | $\delta$ | $C \rightarrow I$ if mediated by $O$ |
| $\varepsilon$ | 0.002 | $\varepsilon$ | $I \rightarrow C$ if mediated by $I$ |

#### Level 3: proliferation level

The cell cycle time for each cell is individually drawn from a distribution depending on the generation number of the cell. The lifetimes of the cells get shorter for higher generation number and the distribution get narrower. The division times (*T*_div_) is drawn from a Gaussian distribution

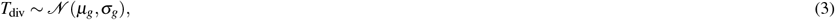

with mean (*μ*_*g*_) and standard deviation (*σ*_*g*_) given per generation in Table 3. These distributions were adapted to experimental data in Olariu *et al*. (2021)^11^.

**Table 3.** Parameter values for proliferation model.

| Generation | $\mu_g$ [h] | $\sigma_g$ [h] |
| --- | --- | --- |
| $g = 0$ | 34 | 13 |
| $g = 1$ | 15 | 5 |
| $g = 2$ | 13 | 5 |
| $g = 3$ | 12 | 4 |
| $g \geq 4$ | 12 | 3 |

When a cell divides, the gene expression levels are copied from the mother cell to the two daughter cells. The states of the Bcl11b regulatory sites are passed on differently depending on the expression level of X. If X *≥* 0, then the regulatory sites are copied from the mother to the daughters. If X = 0, the states are passed on by the following rules: all *C → I, I → I* or *O* with equal probability of each, and all *O → O*.

### Agent-based model

The agent-based model was realised by implementing each cell as an instance of a class. For each cell object, a division time is generated from the distribution given by Eq. (3) and Table 3 and the cell is assigned initial values of gene expression and Bcl11b regulatory site states as outlined in Section Level 3: proliferation level. The rate equations for the transcriptional level (Eq. (1)) are numerically evolved first, followed by evolving the epigenetic rate equations (Eq. (2)) since the epigenetic level depends on the transcriptional level. Each cell object contains a list of all its ancestors and where all its descendants are recorded, respectively.

A simulation of a cell colony is initialised with one single cell with its initial conditions specified in Table 4. A colony simulation was terminated when all cells in a generation had passed 120 hours in simulated time.

**Table 4.** Initial conditions for a simulated T-cell colony.

| Type | Value |
| --- | --- |
| Runx1 | 1 |
| Tcf7 | 2 |
| Gata3 | 1 |
| PU.1 | 5 |
| X | 8 |
| Notch | 7 |
| Closed | 500 |
| Intermediate | 0 |
| Open | 0 |

### Tree plotting

The lineage tree plots for a simulated colony were produced using the Python package ETE 3^26^. The initial cell is placed in the root of the tree. For each cell division, the tree branches in two. The radial length of each branch is proportional to the time span between cell division, i.e. the time-axis points radially outwards. The circular node representing a cell is coloured in black, red or white depending on the cell’s properties. If the cell has an open Bcl11b regulatory region at the time of division, the node is coloured white. If the Bcl11b regulatory region is closed and the level of X at the time of division is above 0, the node is coloured black. If, instead, the level of X is 0, the node is coloured red.

### Last Common Ancestor categories

The definitions of the LCA categories are stated in Figs. 2b and 2c. The precise way of how the classification of each cell in a colony is implemented is summarised in the flowchart shown in Fig. 6.

**Figure 6.**
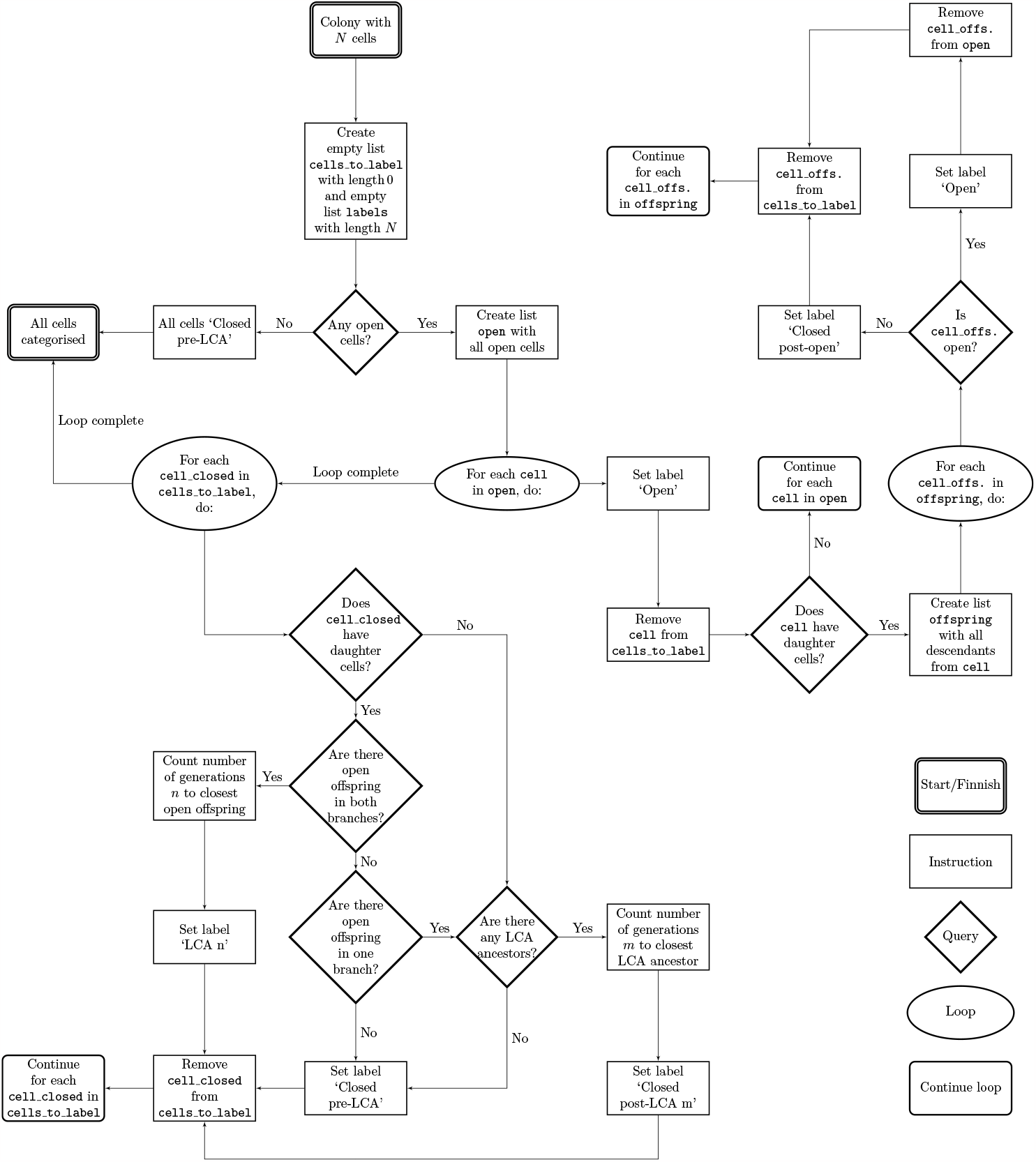
Last common ancestor flowchart. Flowchart over the classification process of the last common ancestor (LCA) categories which starts at the top in the centre.

### Knockdown simulations

Knockdown was implemented by multiplying the production rate of a gene by a knockdown factor. We performed two kinds of knockdowns: initial knockdown and delayed knockdown. When initiating knockdown from the start of the simulation, the initial level of the KD gene was also multiplied by the same knockdown factor. When simulating delayed knockdown, the simulations were carried out the standard way up until the point where the knockdown was initiated, i.e. after 44, 60, 76, 92, or 108 hours in simulated time.

## Acknowledgements

The authors thank Dr. Wen Zhou and Dr. Mary A. Yui for helpful discussions. The authors gratefully acknowledge the support of the United States Public Health Service (NIH) (USPHS grant R01HL119102 to E.V.R. and C.P.)

This work was partially supported by the Wallenberg AI, Autonomous Systems and Software Program (WASP) funded by the Knut and Alice Wallenberg Foundation.

## Author contributions statement

E.A, V.O, E.V.R and C.P. designed the study. E.A. implemented the agent-based multi-scale model and performed all the analysis. E.A. and V.O. wrote the manuscript. All authors provided inputs and comments on the manuscript.

## Additional information

### Data availability

No new data was produced in this study.

### Code availability

Original code is available at https://github.com/Emil-cbbp/agent-based_multi-scale_model.git.

