## Supplementary information for "T-cell commitment inheritance – an agent-based multi-scale model"

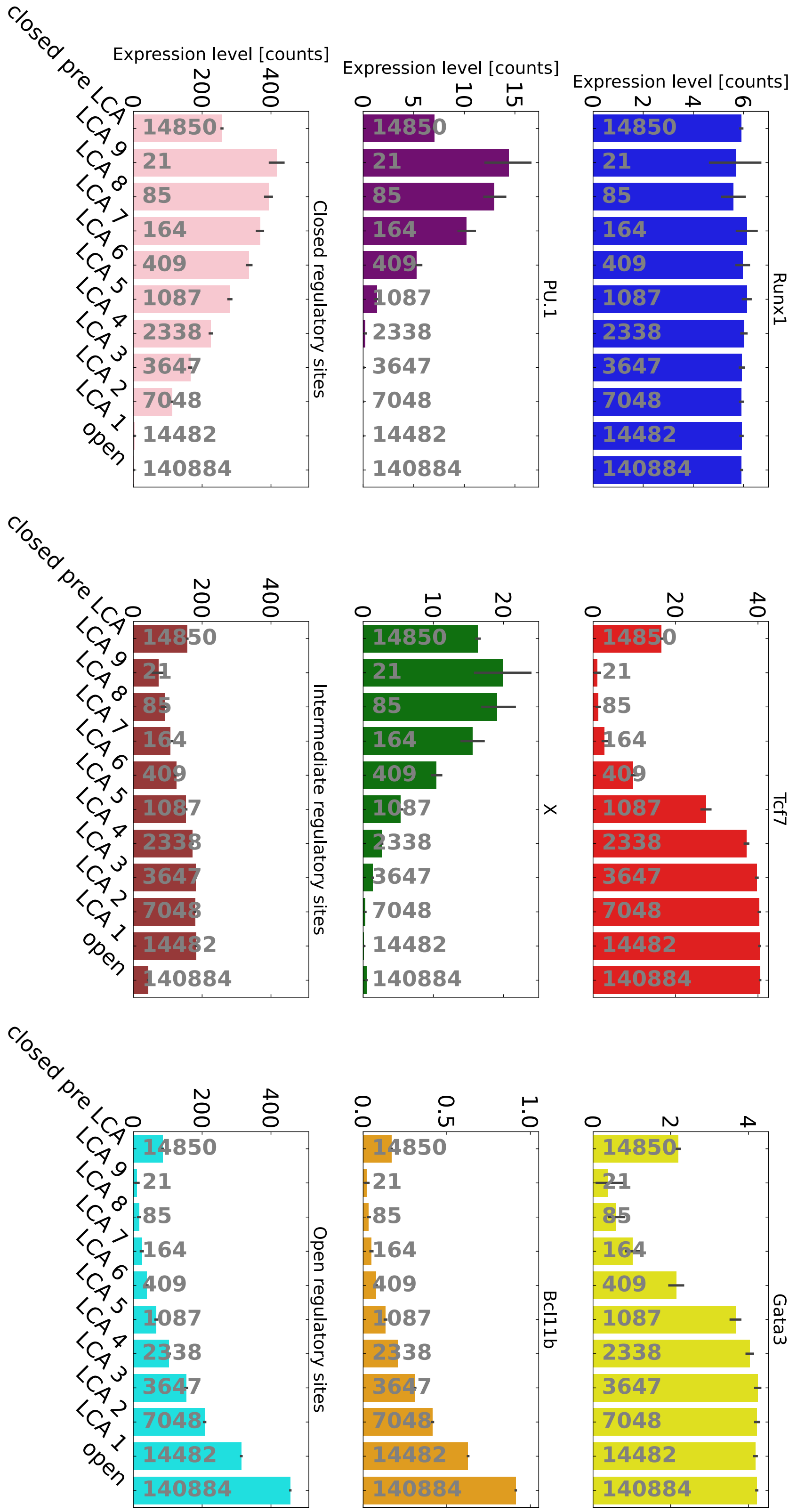


**Supplementary Figure 1: Last common ancestor (LCA) statistics.** Mean expression level for all simulated genes and number or regulatory site statuses for cells belonging to the different LCA categories. The categories are ordered in an approximate developmental order. The grey numbers indicate the number of cells belonging to each category. The coloured arrows highlight interesting events described in the main text. The error bars represent standard deviations.

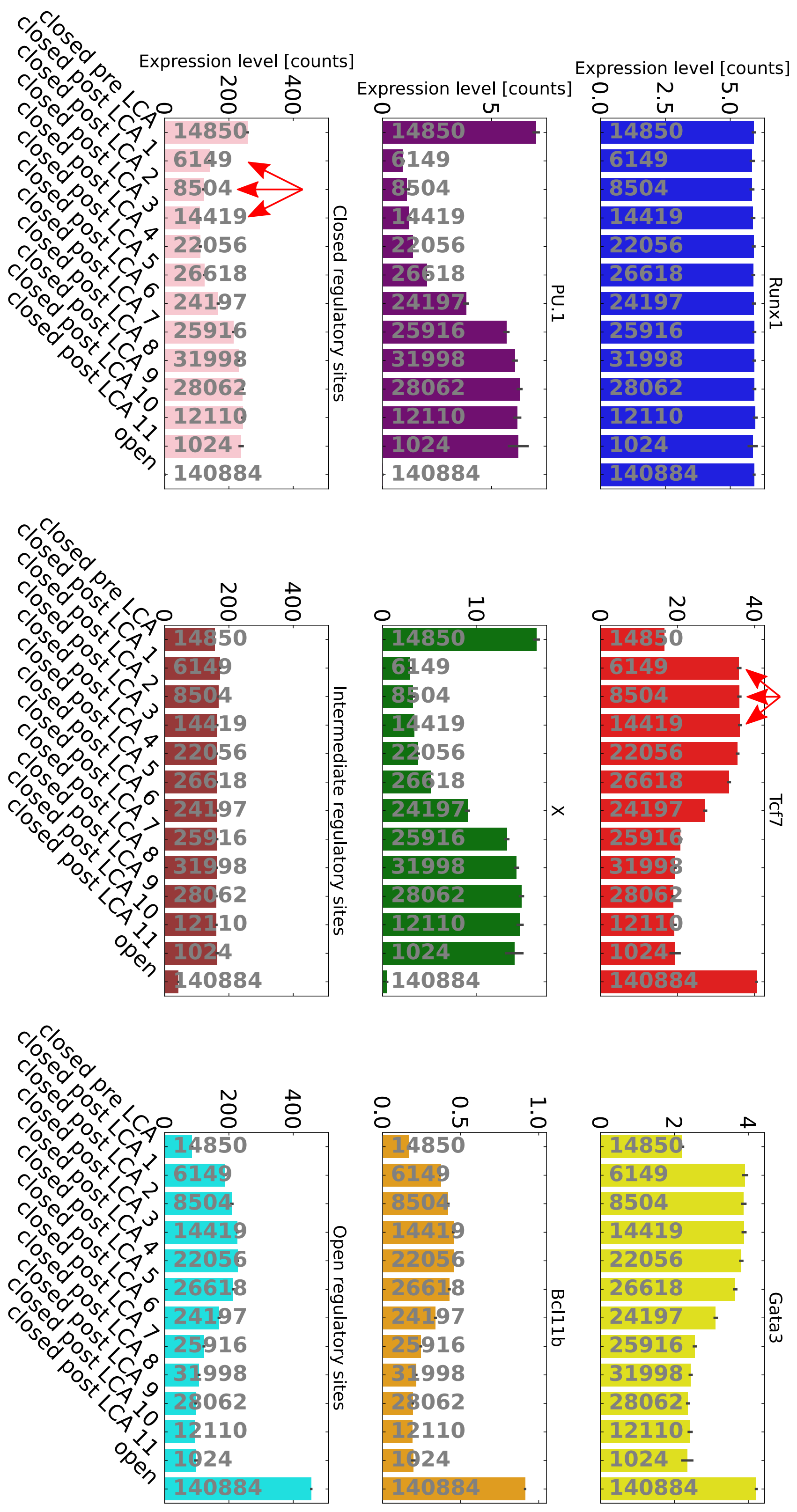


**Supplementary Figure 2: Last common ancestor (LCA) statistics.** Mean expression level for all simulated genes and number or regulatory site statuses for cells belonging to the different 'Closed post-LCA' categories. The categories are ordered in an approximate developmental order. The grey numbers indicate the number of cells belonging to each category. The coloured arrows highlight interesting events described in the main text. The error bars represent standard deviations.

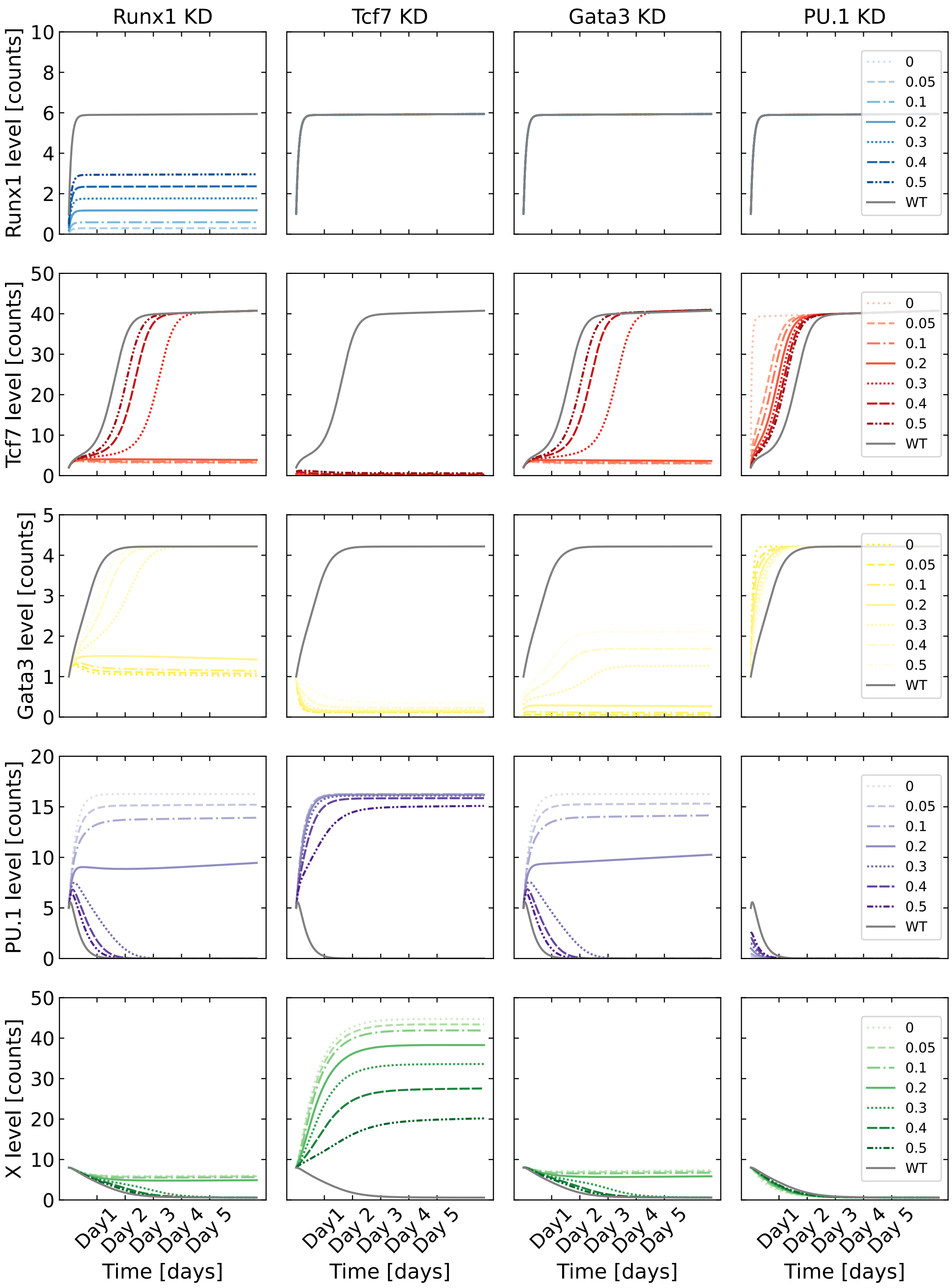


**Supplementary Figure 3: Deterministic knockdown simulations.** Each row shows a deterministically simulated gene where a different gene has been knocked down in each column. The knockdown fraction vary between 5 % and 50 % and is denoted by different line stiles and colour shades. The solid grey line represent wild-type simulations.

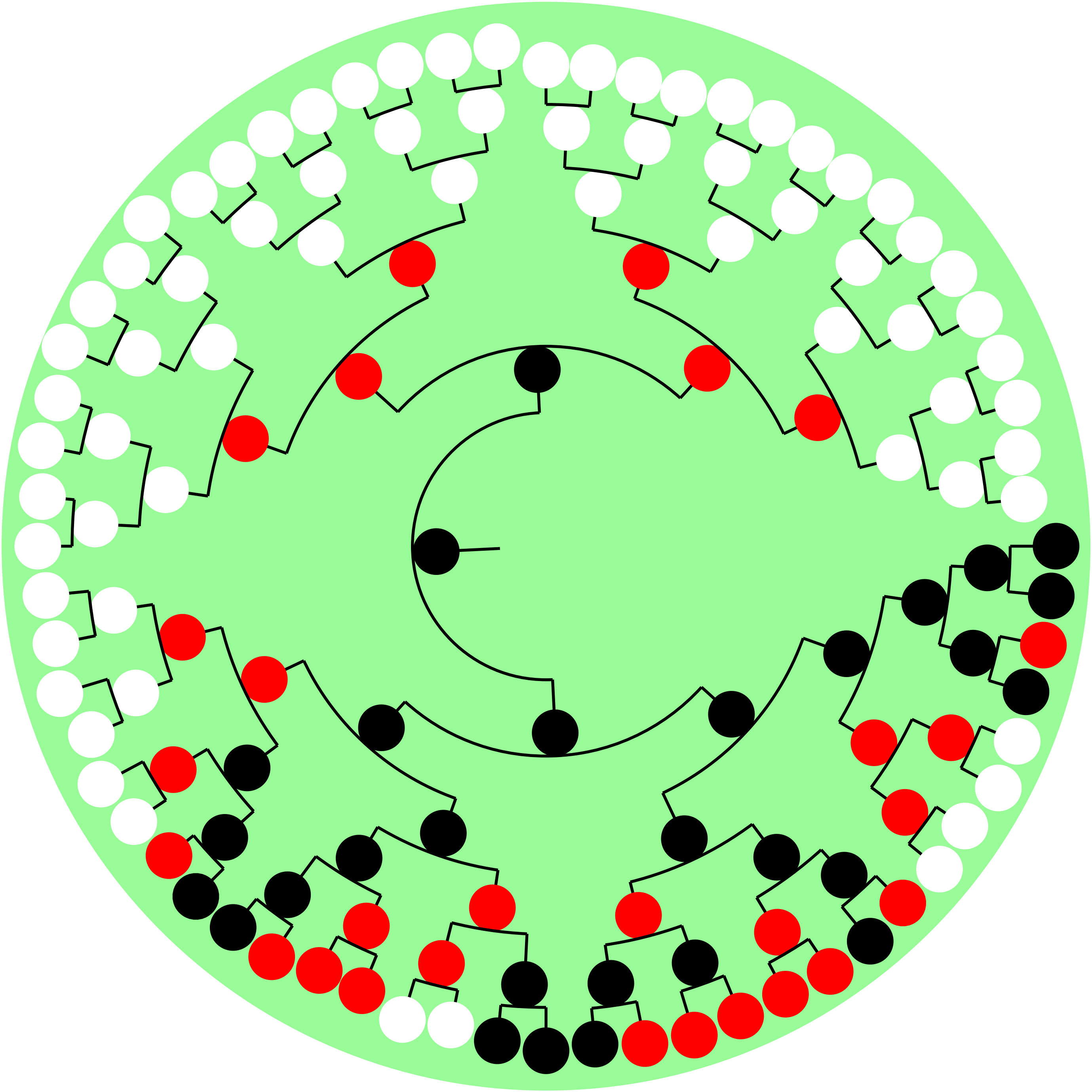

**Supplementary Figure 4: Example lineage tree of simulated T-cell colony with 6 divisions.**

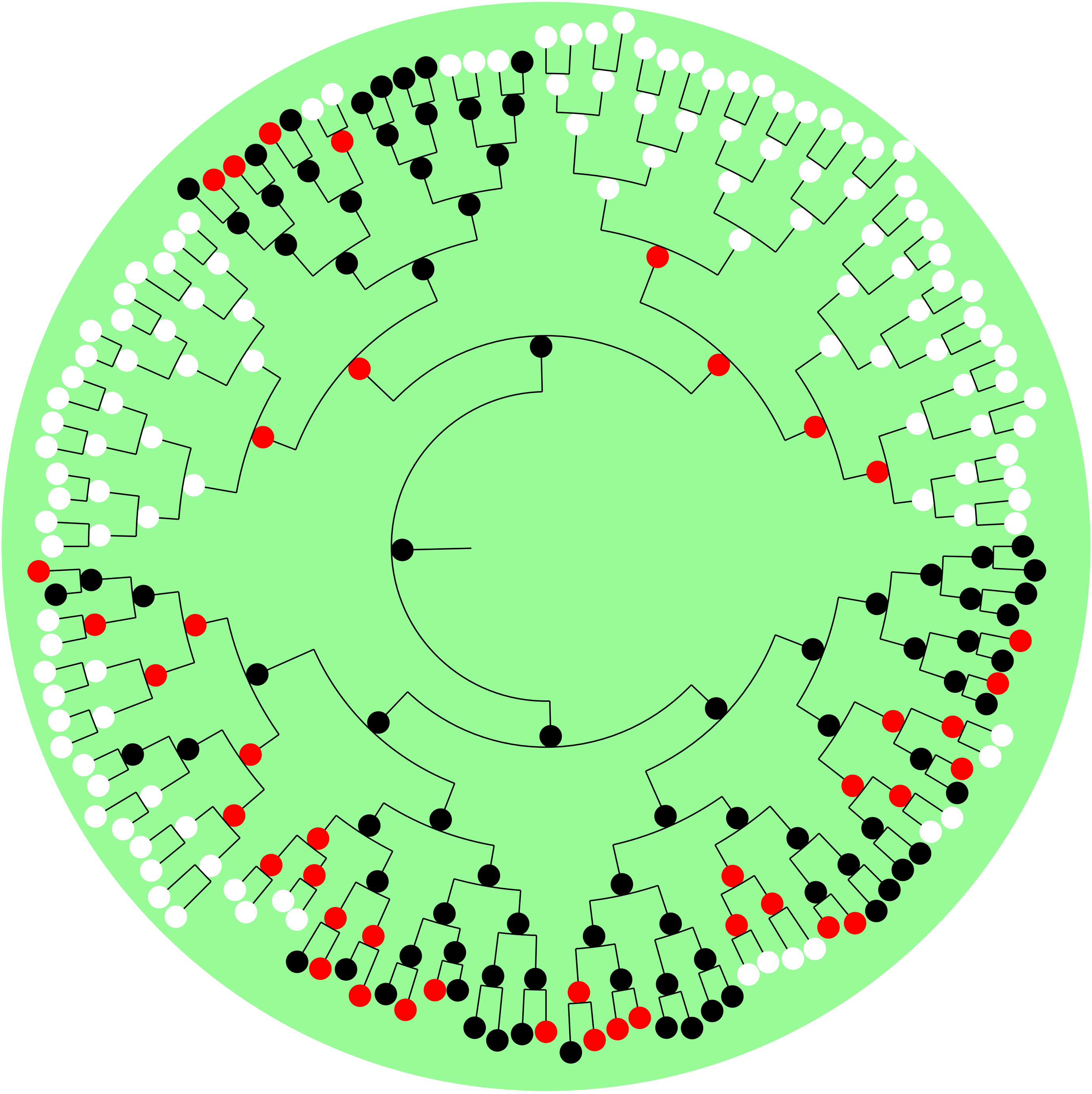

**Supplementary Figure 5: Example lineage tree of simulated T-cell colony with 7 divisions.**

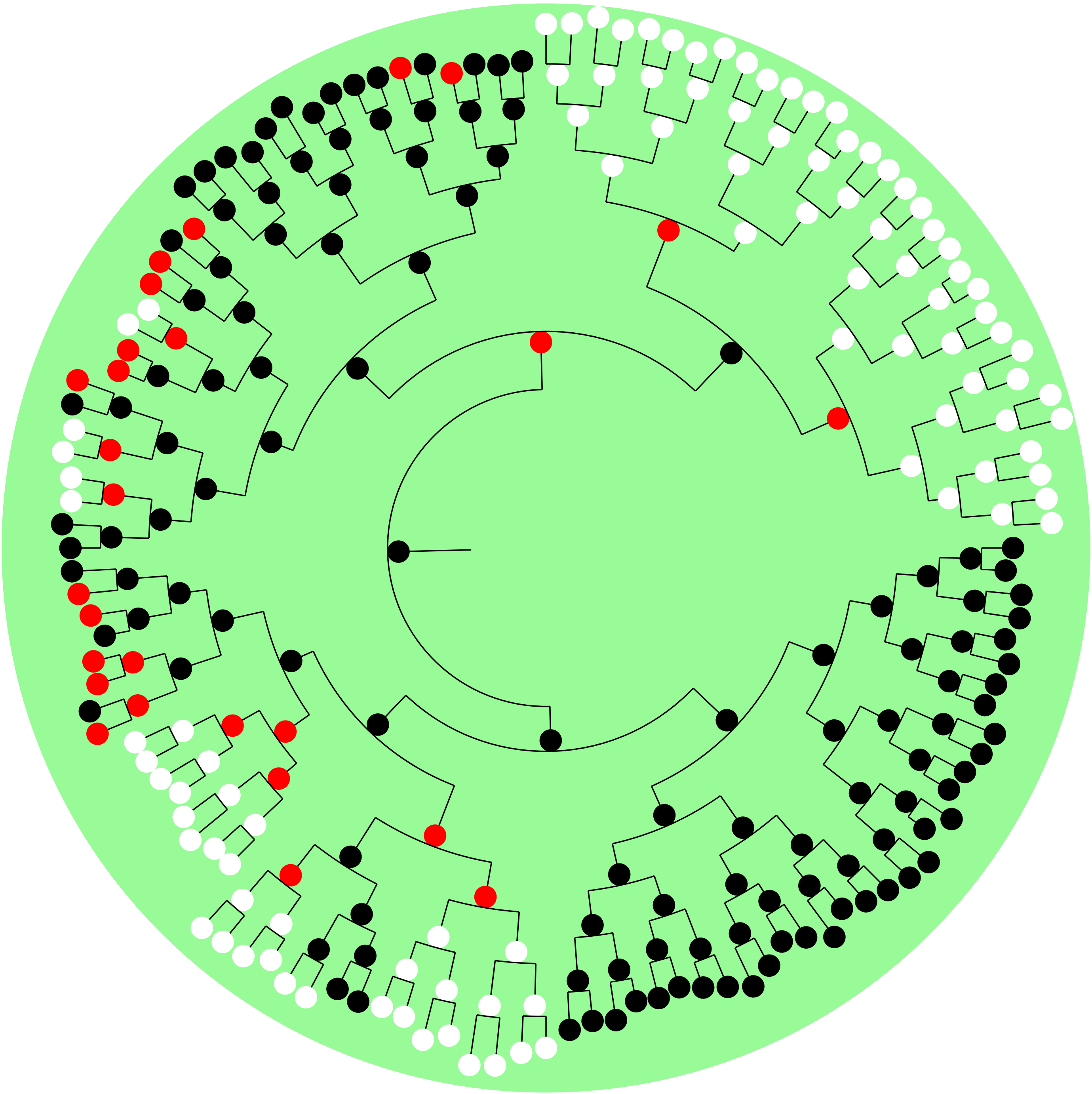

**Supplementary Figure 6: Example lineage tree of simulated T-cell colony with 7 divisions.**

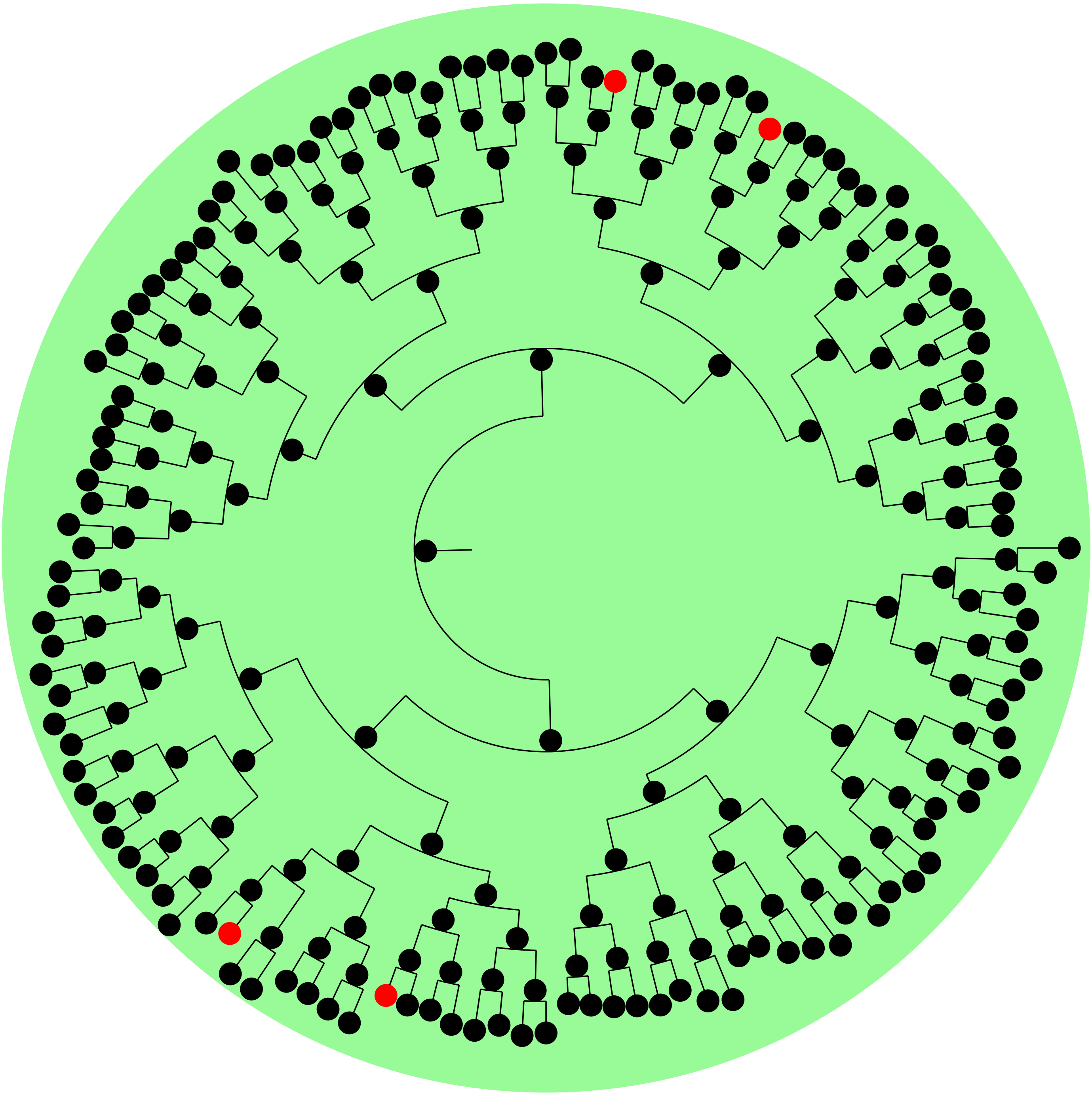

**Supplementary Figure 7: Example lineage tree of simulated T-cell colony with 7 divisions.**

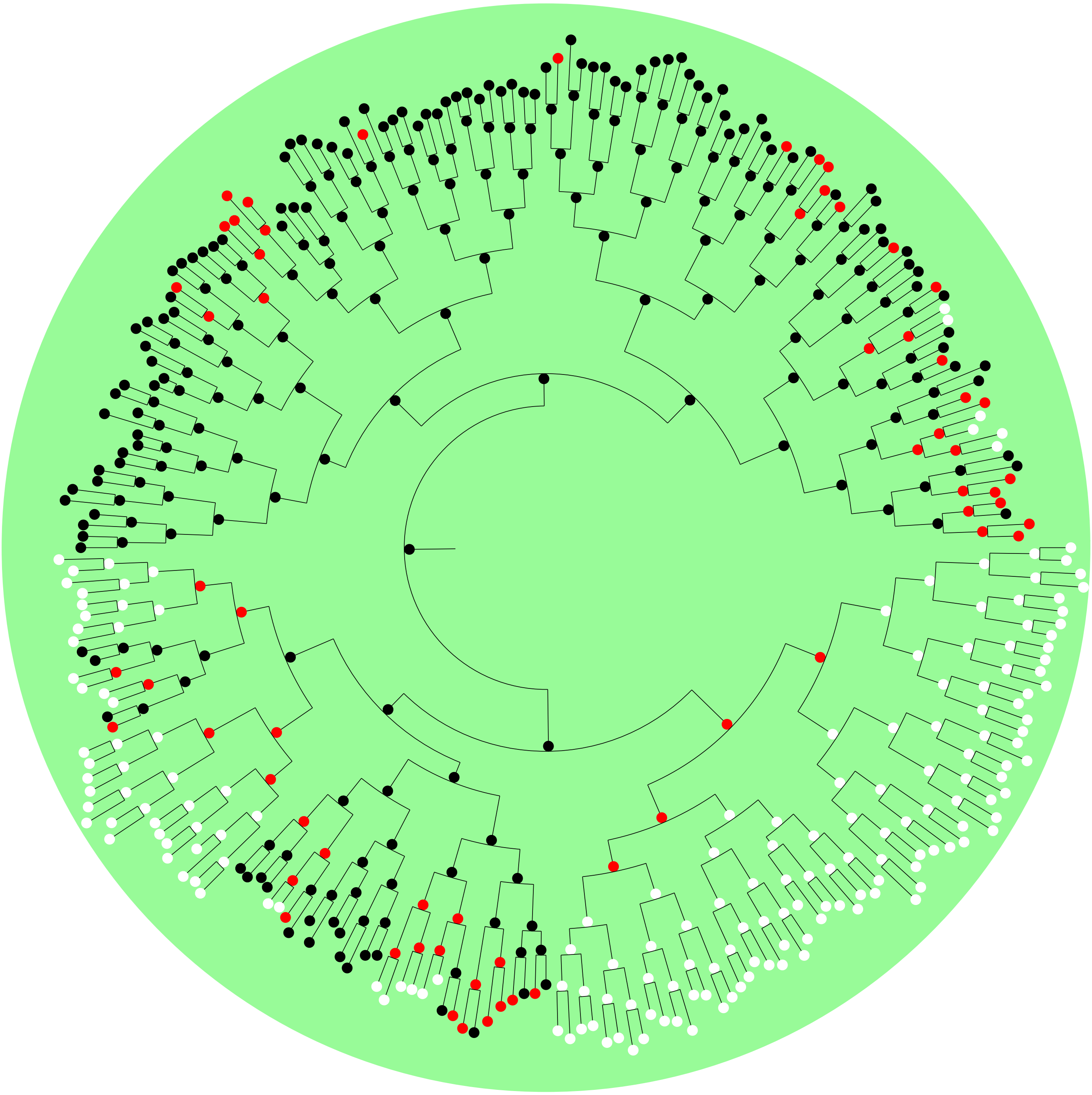

**Supplementary Figure 8: Example lineage tree of simulated T-cell colony with 8 divisions.**

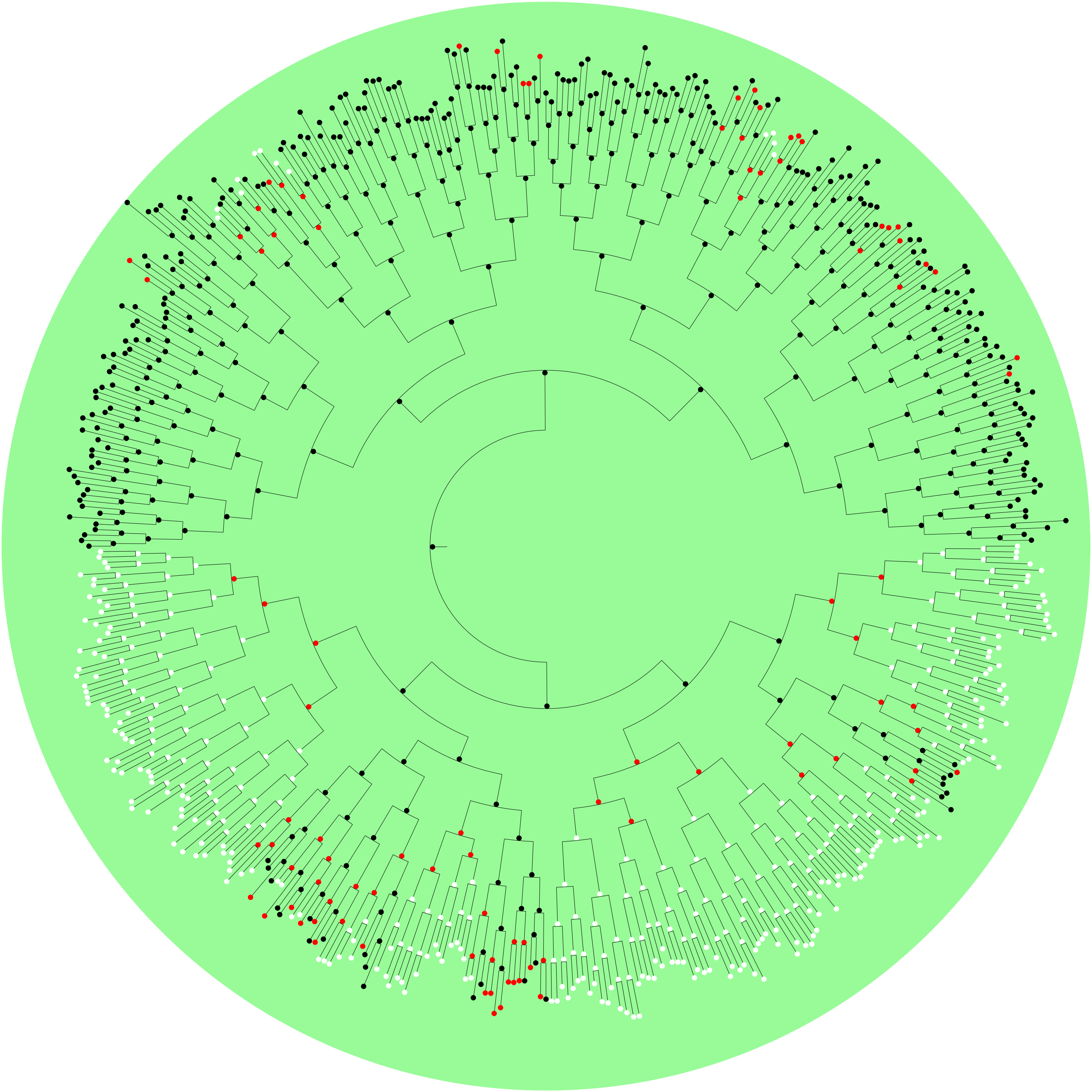

**Supplementary Figure 9: Example lineage tree of simulated T-cell colony with 9 divisions.**
